# From Beeps to Streets: Unveiling Sensory Input and Relevance Across Auditory Contexts

**DOI:** 10.1101/2024.10.17.618668

**Authors:** Silvia Korte, Manuela Jaeger, Marc Rosenkranz, Martin G. Bleichner

## Abstract

This study investigates the neural basis of sound perception in everyday life using EEG data recorded in an office-like environment over 3.5 hours. We investigated how contextual factors such as personal relevance, task complexity and stimulus properties influence auditory processing in ecologically valid settings. By systematically increasing the complexity of acoustic scenes and tasks, we analysed changes in neural responses, particularly in the N100 and P300 components. Our results show that while the P300 is a stable marker of attention in both isolated sounds and complex soundscapes, the N100 is more sensitive to task complexity and environmental factors. This highlights the importance of context in shaping auditory perception. Furthermore, our results suggest that laboratory-based findings can be partially generalised to real-world settings, although task demands significantly influence neural markers. These findings provide new opportunities to study sound perception in naturalistic settings, without sacrificing the control typically afforded by laboratory studies.

## 1 Introduction

Complaints about noise pollution have been around for as long as people have lived together in cities. Its impact on human health and well-being is an increasingly recognized and discussed social problem that has been investigated and proven in a variety of studies (World Health Organization, 2011). It has been shown that habituation to noise varies greatly between individuals and is rarely complete (Basner, Müller, & Elmenhorst, 2011). Heinonen-Guzejev (2008) estimates that about 20 to 43 % of the general population suffers from heightened sensitivity to noise, placing them at a higher risk for negative health outcomes from noise exposure. Studies investigating the adverse effects of noise (defined as unwanted sound) on human health generally refer to the physical properties of noise, which are measurable as loudness, frequency, or sound pressure level (Basner et al., 2014). While much of the research has focused on loud noise and its direct effects, such as hearing loss, non-auditory health impacts can also result from “quiet noise” — sounds that, while not physically loud, are perceived as annoying and stressful over time (Mehrotra et al., 2024; Peris et al., 2020). This form of noise stress has been linked to cognitive deficits, sleep disturbances, depression, and increased suicide rates (Basner et al., 2011; Hygge, Evans, & Bullinger, 2002; Yoon, Won, Lee, Jung, & Roh, 2014).

While the physical impacts of noise on health are well-documented, the question of what constitutes noise is not only highly individual, but also highly complex. While certain physical characteristics of sound, such as high volume or specific frequencies, are generally perceived as disturbing (Rimskaya-Korsakova, Pyatakov, & Shulyapov, 2022), these factors alone cannot fully explain why a sound is considered noise by an individual. In everyday life, we are surrounded by a constantly changing soundscape, and the significance we attribute to these sounds varies depending on the context. Sounds that are relevant to us in one situation may be meaningless or even distracting in another. While the negative health effects of loud noise are well documented, understanding the subjective perception of noise is a significant challenge, that cannot be sufficiently explained by description of the physical properties of sound (Rimskaya-Korsakova et al., 2022).

The complexity of sound perception highlights the need to look beyond physical properties and consider contextual and intrapersonal factors (Hong & Jeon, 2015; Yong Jeon, Jik Lee, Young Hong, & Cabrera, 2011). Emotional state, personal relevance, current task, activity level and intentions all influence how sound is perceived and evaluated (Asutay & Västfjäll, 2012; Debnath & Wetzel, 2022; Rosenkranz, Cetin, Uslar, & Bleichner, 2023; Schlossmacher, Dellert, Bruchmann, & Straube, 2021; Shinn-Cunningham & Best, 2008; Siegel & Stefanucci, 2011). Sounds deemed personally relevant, such as one’s own ringtone or name, command more attention than irrelevant sounds, independent of their physical characteristics (Holtze, Jaeger, Debener, Adilŏglu, & Mirkovic, 2021; Polich, 2007; Roye, Jacobsen, & Schröger, 2013). Moreover, the context in which sound is perceived influences how it is processed (Debnath & Wetzel, 2022). In our study, we therefore pose the question of how personal relevance of a soundscape influences individual sound perception.

One commonly used approach in studying the perception of noise is based on surveys. Although qualitative surveys provide valuable insights into sound perception by accounting for contextual factors, they are prone to cognitive biases, such as priming, false memory, or the peak-end effect (Fredrickson & Kahneman, 1993; Hjortskov, 2017; O’Connell & Greene, 2017). While various studies have attempted to quantify noise perception over time and across environments (see Kjellberg, Landström, Tesarz, Söderberg, and Akerlund 1996; Paunović, Jakovljević, and Belojević 2009; Yoon et al. 2014), no single model has successfully captured the full complexity of what determines sound to be perceived as noise (Pierrette et al., 2012) with the subsequent potential of negative health outcomes. Multiple factors explain variance in noise ratings, including demographic factors such as gender, age and education level, as well as contextual characteristics like perceived acceptability of the noise source, noise expectation and visibility of the noise source (for a detailed review, see Pierrette et al. 2012).

However, surveys offer limited insight into how individuals perceive their current soundscape, as they have to be administered retrospectively. Even with ambulatory assessment methods, where surveys can be done in real time (i.e., as close as possible to a sound event), the challenge remains, that the process of asking a person to reflect on a sound event, may already alter their perception of it and is related to several further challenges (see Schinkel-Bielefeld et al. 2024 for a recent review).

Given the limitations of subjective survey methods, Electroencephalography (EEG) offers an objective alternative for studying how individuals process and perceive sound in real time. EEG can provide insights into the perception of various sound properties such as sound intensity as well as related cognitive processes like attentional arousal, and the detection of auditory expectation violations (e.g.: Näätänen, Kujala, and Winkler (2011); Polich (2007)). Hence, EEG is a valuable tool for capturing sound perception, particularly in situations where immediate, unbiased responses are critical. Its usability in workplace settings is significant, especially in roles requiring sustained attention to a primary task, such as in aviation (Dehais, Roy, & Scannella, 2019), public transportation (Sonnleitner et al., 2014), or healthcare (Rosenkranz et al., 2023), where self-reporting is either impractical or may compromise safety.

EEG studies of sound perception have typically focused on the brain’s response to isolated auditory stimuli over short periods, conducted in controlled laboratory settings that allow for only limited behavioral variation (Lorenzi et al., 2023; Maselli et al., 2023; Nastase, Goldstein, & Hasson, 2020). While these experiments provide valuable insights into basic auditory processing, such as differences in P200 amplitude between noise-sensitive and non-noise-sensitive individuals when exposed to noise (Shepherd, Lodhia, & Hautus, 2019) or how salient events suppress neural responses to ambient sound (Huang & Elhilali, 2020), they may not fully capture the complexity of sound perception in real-world situations. Although there is increasing interest in studying more complex auditory stimuli (e.g.: Ding and Simon 2012; Fuglsang, Dau, and Hjortkjær 2017; Holtze et al. 2021; Horton, Srinivasan, and D’Zmura 2014; Jaeger, Mirkovic, Bleichner, and Debener 2020; Mirkovic, Debener, Jaeger, and De Vos 2015; O’Sullivan et al. 2015; Rosenkranz et al. 2023; Schlossmacher et al. 2021), these controlled experiments still differ significantly from real-world experiences, where individuals encounter a rich variety of sounds, constant environmental changes, and prolonged exposure to auditory stimuli throughout the day (cf. Hasson and Honey (2012)). This raises the question of whether neural processing changes in response to naturally occurring stimuli, which might be perceived as disturbing regardless of their physical properties. The current study, therefore, aims to explore a further question: how does the neural response to auditory stimuli differ between isolated sounds and sounds embedded within a complex soundscape?

EEG studies beyond the lab are becoming increasingly popular and open up a novel approach to study the human brain and behavior (e.g.: Gramann, Ferris, Gwin, and Makeig 2014; Jacobsen, Blum, Witt, and Debener 2021; Ladouce, Donaldson, Dudchenko, and Ietswaart 2019; Reiser, Wascher, Rinkenauer, and Arnau 2021; Scanlon, Jacobsen, Maack, and Debener 2022; Zink, Hunyadi, Huffel, and Vos 2016). However, only few studies have ventured beyond the lab to investigate sound processing in naturalistic environments with even fewer studies focusing on naturally occurring sounds. Rosenkranz et al. (2023); Straetmans, Holtze, Debener, Jaeger, and Mirkovic (2021) demonstrated that responses to continuous stimuli can be recorded while participants were engaging in a bodily active task and Hölle and Bleichner (2023); Hölle, Meekes, and Bleichner (2021) showed that smartphone-based ear EEG can measure sound processing over longer periods of more than 4 hours. However, beyond-the-lab studies face several challenges and the complexity of the ever-changing real world cannot be captured by EEG recordings alone but require environmental context information to account for sources of variance that are not neurally driven (Krugliak & Clarke, 2022). Also, the neural data itself leads to further challenges, such as movement artifacts (Gramann et al., 2014; Jacobsen et al., 2021) and difficulties in interpreting EEG data due to the presence of unknown artifacts (Mathias & Bensalem-Owen, 2019). Additionally, these studies must contend with lower signal-to-noise ratios, and greater environmental variability, which complicates experimental design (Ladouce et al., 2019).

While both traditional lab-based studies and beyond-the-lab studies provide critical insights, each comes with its limitations. Laboratory studies offer the advantage of precise control and measurement but lack ecological validity. In contrast, beyond-the-lab studies capture the complexity of real-world environments, but they face challenges such as movement artifacts and reduced experimental control, complicating the interpretation of EEG data.

Our study seeks to bridge these two approaches by adopting a middle stance. While remaining in a lab-based setting, we simulate a more naturalistic environment, allowing participants greater freedom in a realistic office-work-like scenario. By interacting with complex soundscapes and performing tasks with fewer constraints, participants experience a more flexible and dynamic environment, while we retain enough control to maintain experimental rigor.

Positioned between these two extremes, our study explores whether neural patterns observed in controlled lab conditions can be generalized to more naturalistic yet semi-structured environments. This balanced approach allows us to contribute to a more comprehensive understanding of auditory processing, integrating the strengths of both controlled and real-world research.

In our study, we aim to address the various questions we have raised, which are:

- How does the manipulation of personal relevance change the perception of a soundscape (in the long term)?
- How does the neural response to auditory stimuli differ between isolated sounds and sounds embedded in a complex soundscape?
- Do results from highly controlled experiments generalize to experiments with more degrees of freedom? In our case: Can we monitor changes in sound perception in a realistic (office) working condition?

Specifically, we seek to deepen our understanding of how complex acoustic scenes are perceived and interpreted over extended periods. We use EEG to record participants over several hours while they are exposed to soundscapes and tasks of varying complexity. Additionally, we manipulate the behavioral relevance of the soundscape. Our goal is to enhance the interpretation of sound processing in real-world environments, contributing to a more comprehensive understanding of how sound perception is influenced by context, personal relevance, and stimulus complexity.

## 2 Methods

### 2.1 Participants

The sample consisted of 23 participants (13 female, 10 male) aged between 21 and 37 years (mean: 25.57, standard deviation [SD]: 3.48). All participants were right-handed, had normal (or corrected-to-normal) vision and had no history of neurological, psychiatric or psychological disorders. To ensure that we only include participants with normal hearing, all participants underwent a hearing screening on a day prior to, but close to, the experiment. Audiometric threshold of at least 20 dB HL were confirmed by pure-tone audiometry at octave frequencies from 250 Hz to 8 kHz Using a SIEMENS Unity 2 Audiometer in a sound-proof cabin and with overear headphones. We screened a total of 30 participants of which 23 were eligible.

Prior to EEG recording, participants filled out the Weinstein noise sensitivity scale. The average score was at 3.366 (SD = 0.517), whereas the norm score for this inventory is 3.037 (SD = 0.572). The inventory revealed that six individuals exhibited values of at least one standard deviation above the norm score, indicating heightened sensitivity to noise. Conversely, one individual demonstrated a value below one standard deviation of the norm score, suggesting a diminished propensity for noise sensitivity relative to the average.

Participants gave written informed consent prior to the study and received monetary compensation. The study was approved by the Ethics Committee of the University of Oldenburg.

### 2.2 Procedure

The study examined the neural response to sounds, as measured by EEG. The entire study lasted approximately 6 hours (including breaks, preparation, briefing and debriefing) to provide insight into changes in sound processing over a longer period of time. On the day of the EEG recording, participants were asked to come in with washed hair, to improve data quality. Upon arrival, the participant was taken to the laboratory and any questions about the experimental procedure were answered by the experimenter. We then asked the participant to complete two questionnaires regarding noise sensitivity and general state (described below). After that, we placed the EEG cap. After a short calibration block, we started the experiment, which consisted of six consecutive blocks of 15 to 45 minutes, resulting in a total recording time of 3.5 hours. After each block, the participant could take a break of a self-chosen duration and received instructions for the subsequent block.

#### 2.2.1 Questionnaires: WNSS and General State

Prior to EEG data acquisition participants completed the Weinstein Noise Sensitivity Scale (Weinstein, 1978), a 21-item inventory that asks participants to indicate, on a 6-point Likert scale ranging from “strongly disagree” to “strongly agree” how much they agree with statements related to noise (such as “I wouldn’t mind living on a noisy street if the apartment I had was nice.”). In addition, they completed a questionnaire about their general current state (e.g., hours of sleep, last meal, medications etc.) to ensure eligibility for the study.

### 2.3 Paradigm

All parts of the paradigm (except for the transcription task) were presented using the Psychophysics Toolbox extension (Brainard 1997; Kleiner, Brainard, and Pelli 2007; Pelli 1997, version: 3) on MATLAB 2021b.

#### 2.3.1 General Structure of Block Sequence

There were 6 consecutive blocks, in which the sounds and an additional task became progressively more naturalistic and complex. The design of the blocks was chosen so that results could be compared between them, since only one aspect changed between two consecutive blocks: auditory stimuli, non-auditory task or listening mode.

#### 2.3.2 Experimental Phases

The overall idea was to obtain measures of neural sound processing in three phases and under conditions of varying naturalness. The first phase involved passive listening, the second phase focused on active listening, and the final phase returned to passive listening.

First, we were interested in how task irrelevant background sounds are processed. For this we measured brain activity free of any auditory task to artificial and natural sounds and under varying naturalness of the additional task. In this phase, participants were instructed that the soundscape was not particularly relevant and that they could ignore it. This allowed us to compare neural processing under different levels of stimulus naturalness and the influence of task context.

Second, we manipulated participants’ focus on the sounds by up-modulating the relevance of specific sounds, i.e., a change in pitch in blocks with isolated stimuli and the occurrence of a church bell in blocks with ambient sound. In this phase, participants were asked to respond to these sounds by pressing a key on the keyboard, which required active listening. This allowed us to compare the neural response (quantified as the amplitude of the N100 and P300 component of the ERP) of passive versus active listening for the different auditory materials and the non-auditory tasks.

Third, we asked participants to ignore the previously relevant auditory features again to obtain a measure of what we call “attentional wash-out”. Our prediction was that participants’ P300 response would take some time to return to their baseline level. Thus, we predicted that attention would be washed out over time. For a visual representation of the study design, see Figure 1.

**Figure 1:**
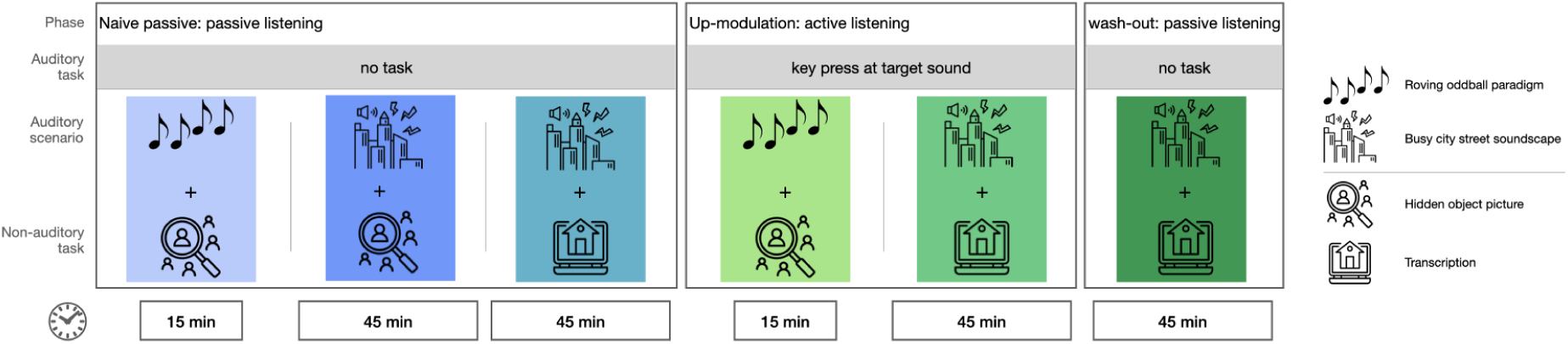
Overview of experimental blocks. Order of the blocks is chosen to ensure naivety concerning target sounds in the naive passive conditions.

#### 2.3.3 Auditory Stimuli

All sound files had a sampling rate of 44.1 kHz. Two auditory scenarios were presented. The first scenario was a roving oddball paradigm (Garrido et al. 2008), based on code publicly available on github.com (Sumner, 2022). In this paradigm 7 stereo pure tone beeps that differed in frequency (500 Hz, 300 Hz, 250 Hz, 200 Hz, 150 Hz, 100 Hz, 50 Hz) were used, with a length of 70 ms and a rise and fall time of 10 ms. For the roving oddball, sequences (trains) of these tones were generated. For each train, a tone at a random pitch was chosen and repeated 5 times, followed by a random addition of 1 to 5 more of the same tone. This resulted in trains ranging from 6 to 10 tones in length. The inter-stimulus interval between tones was 1000 ms. No subsequent train contained the same tones as the preceding train. The roving oddball had a total duration of 15 minutes, resulting in an average of 121 trains. The average volume of the tones was at 55 dB(A).

The second scenario was a pre-recorded soundscape of a busy city street, available on Youtube^1^. It consisted of a variety of ambient sounds, such as streetcars, motorcycles or incomprehensible speech. The street scenario had a total length of 2 hours and 21 minutes, from which we took four segments of 45 minutes each. These segments had a short overlap, since the original sound file was not long enough to cover three non-overlapping hours. The order in which the segments were presented was randomized across participants. Inspired by Brink, Omlin, Müller, Pieren, and Basner 2011, we added a single chime of a church bell (also taken from Youtube^2^) to create an auditory recognition task. The file had a duration of 2890 ms. The church bells were added at 50 random times, with an average distance of 53 seconds between bells. The bells were varied in their spatial position (i.e., they were played from the left, right or both sides). Accordingly, we added bird calls of three different bird species at 50 random time points and with different positions. All bird call files were modified to the same length of 790 ms to facilitate data processing later on. We embedded the church bell and bird calls at a low volume, making them less noticeable compared to the background noise. The sounds were chosen to be natural in the overall soundscape of the busy city. This approach was inspired by the study by Schlossmacher et al. (2021), where words were embedded in noise and were only recognized by the participants when they were made aware of the words. The resulting overall soundscape had an average volume of 51 dB(A).

#### 2.3.4 Non-auditory Task

Participants engaged in one of two non-auditory tasks throughout the experiments depending on the block. The first task was a hidden object picture task involving visual search. A hidden object picture is a static picture with a high level of detail. There are a variety of visual objects and activities and participants had to search for a specific aspect of the picture (e.g.: “Find the black cat”). The task is similar to “Where is Waldo?/Where is Wally?”, the famous children’s game.

The hidden object pictures were comic-style and showed different scenes, such as a library, a playground or a hair salon. In each picture, 40 different objects were marked as targets. Participants were tasked with searching for individual objects, and the target object was written at the top of the screen throughout. Using the mouse, they could click on any object. If they clicked on the target, a message on the screen informed them of the successful identification. If they clicked any other object, they received a message instructing them to continue searching. Participants had unlimited time and unlimited attempts to find the target object. If they could not find a target or were uncertain about a word, they could skip to the next target by pressing the right arrow key on the keyboard. Participants had to search for an object for at least 30 seconds before they could choose to skip an object. If participants identified all 40 objects in a picture before the end of an experimental block, they were presented with a new hidden object picture. Participants were presented with a maximum of 5 hidden object pictures (depending on their individual task speed). The number of targets was deliberately set so high that participants would not be able to find all the targets in the allotted time, to ensure that all blocks were of equal length for all participants. The order of the pictures was randomized across participants.

The second task was a transcription task that resembled simple office work. The task was taken from the citizen science project “world architecture unlocked” on “Zooniverse” and consisted of transcribing handwritten information about architectural photography^3^. The handwritten information had to be matched to given categories, such as city, name of architect, or name of building. The task required no prior knowledge about architecture, yet was challenging enough that participants had to stay focused to complete the task. Participants were encouraged to use online search engines and online maps to correctly match the handwritten information under the photos to the required categories. The combination of deciphering, typing, and searching the internet met the requirement of resembling office work and provided higher task complexity than the hidden-object picture task.

#### 2.3.5 Experimental Blocks

The experimental phases described above were divided into 6 consecutive blocks, as can be seen in Figure 1. The naive passive phase had 3 blocks, the up-modulation phase had 2 blocks and the wash-out phase had one block.

Both auditory scenarios were presented in either passive or active listening mode. In the passive listening mode, participants were informed that there would be background noise but that it was irrelevant, and could be ignored. In the active listening mode, participants were asked to respond to specific acoustic events for each of the auditory scenarios. For the roving oddball, they were asked to indicate a change in pitch from one tone to the next by pressing the F4 key on the keyboard. For the street scenario, they were asked to indicate the occurrence of the church bell by pressing the F4 key. The response key was chosen to avoid a potential conflict in the non-auditory task, where participants also had to use the keyboard. All other sounds, including the bird calls, were not behaviorally relevant and never required a response. They were used to facilitate the ERP analysis, so that the relevant events (i.e., church bells) could be compared with non-relevant events (i.e., bird calls).

In detail, the experimental blocks were as follows:

- Block 1 (Naive passive listening): Participants listened to a 15-minute sequence of the roving oddball while completing the hidden object picture task. They were not required to respond to the sounds.
- Block 2 (Naive passive listening): Participants listened to a 45-minute sequence of the street soundscape while working on the hidden object picture task. They were not required to respond to the sounds.
- Block 3 (Naive passive, passive listening): Participants listened to a 45-minute sequence of the street soundscape while performing the transcription task. They were not required to respond to the sounds.
- Block 4 (Active listening, up-modulation): Identical to Block 1, but now participants were instructed to respond to the first tone of each train.
- Block 5 (Active listening, up-modulation): Identical to Block 3, but now participants were instructed to respond to the church bell sounds.
- Block 6 (Passive Listening, wash-out): Identical to Block 3. Mind that participants were now required to ignore the previously relevant church bell sound again. They were not required to respond to the sounds.

The order of blocks was identical for all participants. This was necessary to assure that participants were naive to the target sounds in the passive listening blocks.

### 2.4 Hypotheses

We analyzed the EEG data for early and late components of the ERP which are the N100 component, known to reflect early auditory processing and stimulus intensity (Näätänen & Picton, 1987) and the P300 component, known to reflect higher cognitive processing, such as attention (Polich, 2007). We postulated the following hypotheses:

- Hypothesis 1: The amplitude of the P300 component to target stimuli will be greater for a target sound in the active listening condition compared to the passive listening condition. This effect has been consistently found in laboratory experiments (e.g.: Polich 2007). To test this hypothesis, we compared the ERPs from block 1 (passive listening to the roving oddball + hidden object picture task) to block 4 (active listening to the roving oddball + hidden object picture task) and from block 3 (passive listening to the street soundscape + transcription task) to block 5 (active listening to the street soundscape + transcription task).
- Hypothesis 2: Processing of irrelevant sounds, as represented by the N100 of the ERP to non-relevant beeps in the roving oddball and non-relevant added bird sounds in the street soundscape, will generally be larger in the active listening condition than in the passive listening condition because participants have to scan the entire soundscape for the target. To test this, we compared the same blocks as in hypothesis 1.
- Hypothesis 3: We hypothesized a difference in P300 amplitude for relevant features of the auditory scene in active versus passive listening between the blocks with more controlled experimental features (i.e., roving oddball + hidden object picture task) and blocks with less controlled experimental features (i.e., street soundscape + transcription task). We predicted that the P300 amplitude would show a higher amplitude in active versus passive listening in blocks with more experimental control than in the blocks with less experimental control. To test this, we compared the grand average between the effects of block 1 (hidden object picture task + passive roving oddball) and block 4 (hidden object picture task + active roving oddball) versus block 3 (transcription task + passive naive street soundscape) and block 5 (transcription task + active street soundscape).
- Hypothesis 4: The amplitude of the N100 component of the ERP will be greater during passive listening to the street soundscape when working on the hidden-object picture task compared to the transcription task. We hypothesized that the transcription task, resembling simple office work, would elicit a smaller N100 component for relevant (bell) as well as irrelevant (bird) sounds in the street soundscape compared to the street soundscape while working on the hidden object picture task. To test this hypothesis, we compared block 2 (passive listening to the street soundscape + hidden object picture task) to block 3 (passive listening to the street soundscape + transcription task).
- Hypothesis 5: We hypothesize a gradual down-modulation of the personal relevance in the street soundscape in the last block (transcription task + wash-out street soundscape), as indicated by a reduction in P300 amplitude over time. We compared the P300 amplitudes of the data in increments of thirds between the passive naive listening condition and the wash-out condition. Our expectation was that there would be a significant difference between each first third but not between each final third.

### 2.5 Data Acquisition

#### 2.5.1 Description of Lab Setup

Participants were seated in a soundproof recording booth at a desk with a screen (Samsung, SyncMaster P2470) in front of them. The auditory material was presented free-field through two loudspeakers (Sirocco S30, Cambridge Audio, London, United Kingdom) positioned at a 45-degree angle to the left and right and at a distance of approximately 0.5 m at ear level. A mouse and a keyboard were placed on the desk in front of the participant which were used to indicate target sounds and for working on the non-auditory tasks. For relevant events in the auditory scenes and additional tasks, a marker was generated using the Lab Streaming Layer (LSL) library^4^. Keyboard input was recorded using key capture software from the LSL library^5^. The Lab Recorder software^6^ was used to ensure the temporal synchronicity of the EEG data, the event markers and the keyboard capture. Files were saved as .xdf and organized using the Brain Imaging Data Structure (BIDS) format (Gorgolewski et al., 2016) with the EEG data extension (Pernet et al., 2019).

#### 2.5.2 EEG system

We used a 24-channel EEG cap (EasyCap GmbH, Hersching Germany) with passive Ag/AgCl electrodes (channel positions: Fp1, Fp2, F7, Fz, F8, FC1, FC2, C3, Cz, C4, T7, T8, CP5, CP5, CP1, CPz, CP2, CP6, TP9, TP10, P3, Pz, P4, O1, O2). The mobile amplifier (SMARTING MOBI, mBrainTrain, Belgrade, Serbia) was attached to the back of the participants’ head in a small pocket of the EEG cap, so that they could move their head freely throughout the experiment (and it was also easy to leave the lab, e.g. for a bathroom or lunch break). In addition, we collected gyroscope data from the amplifier to track the participants’ head movements. The data were transmitted via Bluetooth using a Bluetooth dongle (BlueSoleil) connected to a desktop computer that was also used for stimulus presentation. EEG and gyroscope data were transmitted to LSL via the SMARTING Streamer software (v3.4.3; mBrainTrain, Belgrade, Serbia) and recorded with the lab recorder at a sampling rate of 250 Hz.

#### 2.5.3 Measurement Procedure

After applying the cap, the skin beneath each electrode was cleaned with 70 % alcohol and abrasive gel (Abralyt HiCl, Easycap GmbH, Germany). The electrodes were then filled with abrasive gel to improve the conductance between the skin and the electrodes. Impedances were kept below 10 kΩ at the beginning of data collection and were checked and improved as necessary between the blocks.

Given the length of the experiment, all participants were asked to bring food to allow for an extended lunch break between blocks. The time of the lunch break was flexible, with the only restriction that the break could not be between the last two blocks (active listening and wash-out) so as not to compromise the experimental manipulation.

### 2.6 Data Analysis

All data analysis was carried out using MATLAB 2021b with the EEGLAB toolbox (Delorme and Makeig 2004; version: 2021.1).

#### 2.6.1 EEG Pre-processing

To clean the EEG data, we applied a pre-processing to it. First, we filtered the data between 1 and 40 Hz (default settings of the pop eegfiltnew function). Then we detected bad channels, using the clean artifacts function of EEGLAB with “channel crit maxbad time” at default setting and stored them for later interpolation. We then cut the data into segments of 1 second length. For these segments, we applied an artifact rejection with a probability criterion of +/- 3 SD from the mean. This was done to improve the independent component analysis (ICA) training. Second, we combined all the data obtained from the first cleaning step to compute the weights for an ICA. We used the runica function from EEGLAB and chose the extended training method. The ICA weights were then back-projected onto the raw data of each block separately. For component rejection, we applied the ICLabel algorithm (Pion-Tonachini, Kreutz-Delgado, & Makeig, 2019) and rejected components with a probability *≥* 80 % of being an artifact of any class (i.e.: eye, muscle, heart, other). Additionally, we rejected components based on visual inspection where necessary, as the ICLabel algorithm is only optimally trained on stationary data where participants were constrained in their movements, which does not fully match the characteristics of our setup, where participants were free to move within the scope of the task. After ICA cleaning, we filtered the data (low pass: 0.5 Hz, high pass: 20 Hz), interpolated any bad channels and re-referenced the data to the mean of electrodes Tp9 and Tp10.

#### 2.6.2 ERP Analysis

Prior to epoching, we corrected audio events for a constant delay of 35 ms. EEG data were then epoched around the audio events of interest. For the roving oddball (blocks 1 and 4), we epoched between -200 and 1000 ms relative to the onset of the first tone and last tone of each train. For the street soundscape (blocks 2,3,5 and 6), we created epochs of 2.2 seconds length (-200 to 2000 ms) around each event of a church bell and each event of a bird call. The longer epoch duration in the street soundscape condition accounted for the longer response window, as explained above. We found a time-locked artifact in the active street condition (block 5) in the time range of reaction times that can be explained by the fact that some participants were moving their head down to face the keyboard and find the response key when a target appeared. This artifact is especially pronounced in the frontal electrodes but does not contaminate the time windows of interest for the N100 and P300 analysis.

Epochs for both conditions were baseline corrected (-200 to 0 ms) and epochs that deviated at least 3 SD from the mean were rejected. The remaining epochs were used for the ERP analysis. For analyzing the P300 component, we did a channel selection of channels P3, Pz and P4, since the parietal channels usually show the highest contribution to the P300 peak of the ERP (Polich, 2007). For analyzing the N100 component we selected frontal channels FC1, Fz and FC2 (Näätänen & Picton, 1987).

Table 1 in the supplementary material holds the overview of sample sizes per block after correcting for corrupted or missing data.

**Table 1:** Overview of reasons for excluding participants from the ERP and statistical analyses per block.

| Block | N | Reason for exclusion |
| --- | --- | --- |
| 1 | 22 | 1x bad data quality |
| 2 | 22 | 1x bad data quality |
| 3 | 22 | 1x bad data quality |
| 4 | 21 | 1x missing data,<br>1x bad data quality |
| 5 | 21 | 1x bad performance in auditory task,<br>1x bad data quality and non-compliance in non-auditory task |
| 6 | 21 | 1x bad performance in auditory task in former block,<br>1x bad data quality and non-compliance in non-auditory task |

#### 2.6.3 Behavioral Data

For the auditory task, we calculated hits, misses and false alarms for the target sounds. Additionally, we calculated the response times for the hits to the auditory task in the active listening blocks. For the roving oddball, a hit was defined as a response key press after each first tone of a train within a time range of 200 - 1000ms after tone onset. A false alarm was defined as a response key press after any tone other than the first tone of a train. Similarly, for the street soundscape, a hit was defined as a response key press after the church bell (time range: 200 - 2000ms), while a false alarm was defined as a response key press at any other time or event. The longer response window for the street soundscape was chosen because participants could not rest their hand on the response key while working on the transcription task, as was possible (and encouraged) for the hidden object picture task. Moreover, the longer response window accounted for the increased task difficulty, introduced by the background noise, resulting in an auditory discrimination task instead of an auditory detection task as in the roving oddball condition (Deshpande, Brandt, Debener, & Neher, 2022).

For the hidden object picture task, we looked at the average number of objects correctly identified in 15 minutes and examined whether task performance changed over time (i.e.: participants identified more/less objects per block).

There was no direct behavioral measure for the transcription task. To get an estimate of the overall task engagement of the participant in this task, we calculated the number of keystrokes per minute.

#### 2.6.4 Statistical Testing

Wilcoxon signed-rank test was used for all comparisons in testing the hypotheses and the task performances in the additional tasks. We assumed evidence for an effect at *α* =.05. To control for Type I errors (false positives) arising from multiple comparisons, we applied a Bonferroni correction for multiple testing in our analysis. Given that certain data blocks (especially bell-epochs from block 3) were used across multiple hypotheses, we employed an overarching correction approach. This method ensures that all comparisons involving the same dataset are accounted for across different hypotheses, rather than treating each hypothesis in isolation. For each hypothesis, a unified correction was applied based on the maximum number of comparisons in which the same data block was used across all hypotheses. This led to a correction for 6 comparisons in hypotheses 1, 3, 4 and 5 and 2 comparisons in hypothesis 2 (detailed explanation in supplement). For comparisons of the N100 amplitude, we used the average over channels FC1, Fz and FC2. For comparisons of the the P300 amplitude, we used the average over channels P3, Pz and P4. In blocks, where the roving oddball was applied, the time window for the N100 was set to 50 - 160 ms and the time window for the P300 was set to 350 - 800 ms. In case of the street soundscape we used a time window of 90 - 200 ms for the N100 analysis and a time window of 450 - 900 ms for the P300 analysis. Time windows were averaged across their whole length.

## 3 Results

### 3.1 Auditory Task Performance

The average hit rate in the active roving oddball was at 81.56 % (SD = 20.79). The average hit rate for the active street soundscape was comparable at 86.52 % (SD = 19.50). In the roving oddball, there was a total of 120 tone changes (i.e., 120 targets). In the street soundscape were 50 occurrences of the bell sound (target). Although the numbers are comparable, it can be seen in figure 2a, that there was a higher variance in the performance in the roving oddball.

**Figure 2:**
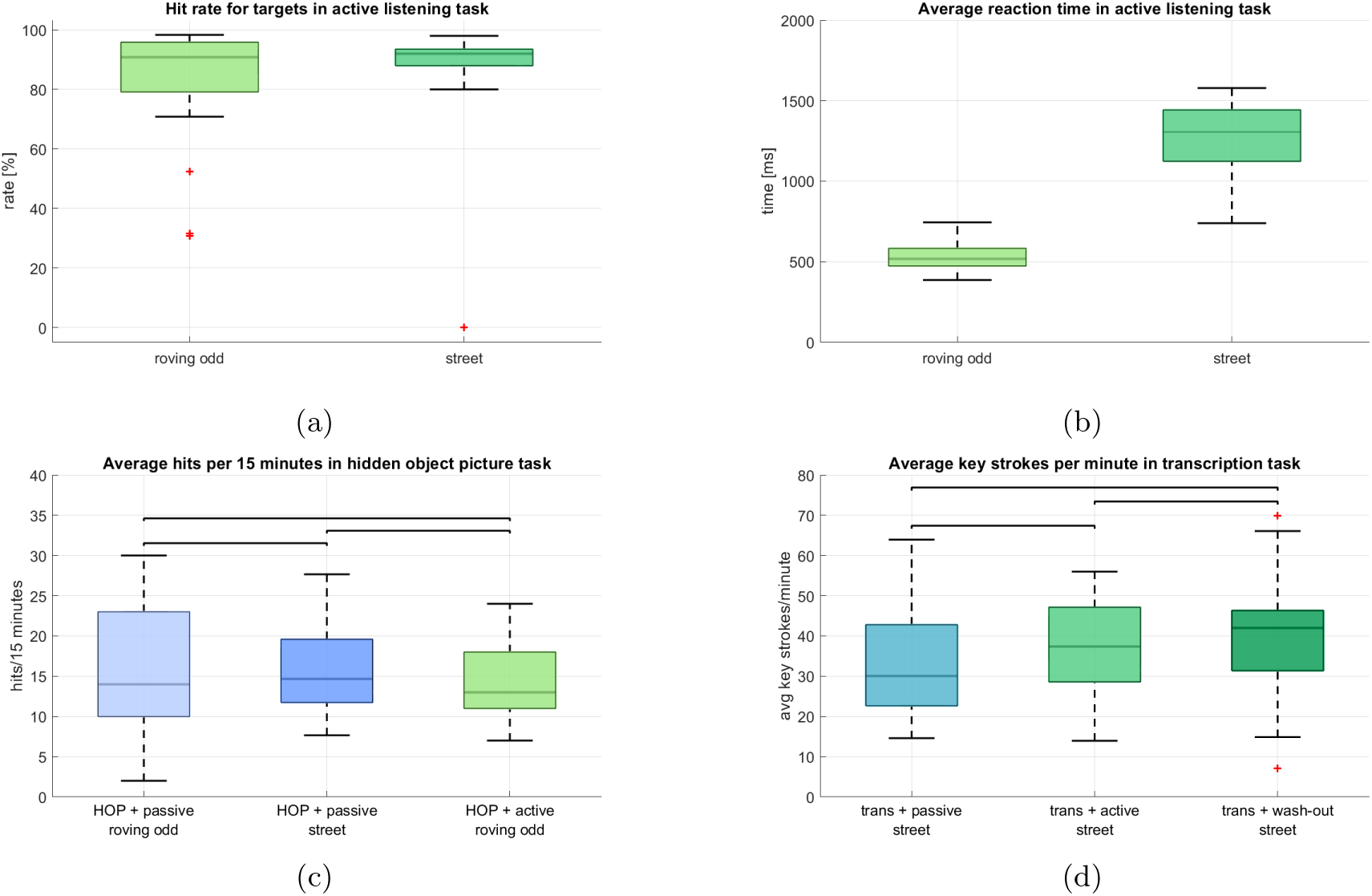
Behavioral task performances in the auditory and non-auditory tasks. **a** Comparison of the hit rate in the auditory tasks in active listening. **b** Comparison of reaction times in the auditory tasks in active listening. **c** Comparison of the hit rates for the 3 blocks in which the hidden-object picture task was performed. Data is displayed as a 15-minute-average to grant comparability between blocks. **d** Comparison of the average absolute number of key strokes per minute in the transcription task.

The average reaction time in the roving oddball was at 525.79 ms (SD = 82.01) and 1258.10 ms (SD = 235.50) in the street soundscape. The large difference can partly be explained by the fact that for the roving oddball participants could rest their left hand on the target key, such that they could immediately press it when they heard a target. In the street soundscape condition, participants were asked to press the same key but they were not able to rest their hand on it, since they needed both hands for typing during the transcription task. It can be seen in figure 2b that the roving oddball yielded a smaller variance in reaction times, while there was a larger variance in the street soundscape condition.

### 3.2 Additional Task Performance

Figure 2c and 2d show the performance in the additional tasks (i.e., hidden object picture and transcription). For the hidden object picture task (2c), we display the average number of correctly identified targets in 15 minutes to ensure comparability between the blocks, since they differed in total length (15 minutes vs 45 minutes). Under passive naive listening of the roving oddball (block 2), participants on average correctly identified 16.35 targets (SD = 8.29). Under passive naive listening of the street soundscape they had an average performance of 15.80 targets (SD = 5.69) and in the active listening block they identified on average 14.14 targets (SD = 4.99).

To obtain a behavioral measure for the transcription task, we counted the average number of keyboard strokes per minute and participant for each block (2d). Under passive naive listening of the street soundscape (block 3), participants had on average 34.23 key strokes per minute (SD = 14.78). In the active listening condition (block 5) they had an average of 37.85 (SD = 13.09) and in the wash-out out condition (block 6) they had an average of 40.40 (SD = 15.37), It can be seen that in both tasks there is no clear pattern of performance gain or loss over time as could have been expected especially for the transcription task, where participants had to learn the task initially. We tested this using Wilcoxon signed-rank test and corrected for 3 comparisons to investigate the group differences for each block of the hidden object picture task and the transcription task separately. There were no significant differences between the blocks for the performance in the hidden object picture task (HOP + passive roving odd vs. HOP + passive street: adj. p = 0.945, Z = 0.069; HOP + passive roving odd vs. HOP + active roving odd: p = 0.088, Z = 1.707; HOP + passive street vs. HOP + active roving odd: p = 0.158, Z = 1.213). Also, there were no significant differences between the blocks for the transcription task (trans + passive street vs. trans + active street: p = 0.128, Z = -1.521; trans + passive street vs. trans + wash-out street: p = 0.039, Z = -2.100; trans + active street vs. trans + wash-out street: p = 0.189, Z = -1.315).

### 3.3 Hypotheses Testing

For Hypothesis 1 we expected a larger P300 amplitude to target stimuli (i.e., first tones in roving oddball and bell sounds in street soundscape) in the active listening condition as compared to the passive listening condition. We compared the respective blocks for the roving oddball (i.e., block 1 and 4) and the street soundscape (i.e., block 3 and 5) separately and performed a Wilcoxon signed-rank test for each comparison. We found significant differences between the active listening and the passive listening condition for the first tones in the roving oddball (adj. p *≤* 0.001, Z = -10.192, adjusted for 6 comparisons, Figure 3a) as well as for the bell sounds in the street soundscape (adj. p *≤* 0.001, Z = -10.192, adjusted for 6 comparisons, Figure 3b), as displayed in Figure 3. The mean amplitudes in the time window of interest for the roving oddball were m = -0.402 (SD = 0.415) in the passive listening condition and m = 2.631 (SD = 0.542) in the active listening condition. For the street soundscape, we found a mean amplitude of m = -0.035 (SD = 0.222) in the passive listening condition and m = 3.294 (SD = 1.252) in the active listening condition. We found evidence for the first hypothesis being in line with existing literature (e.g.: Polich 2007).

**Figure 3:**
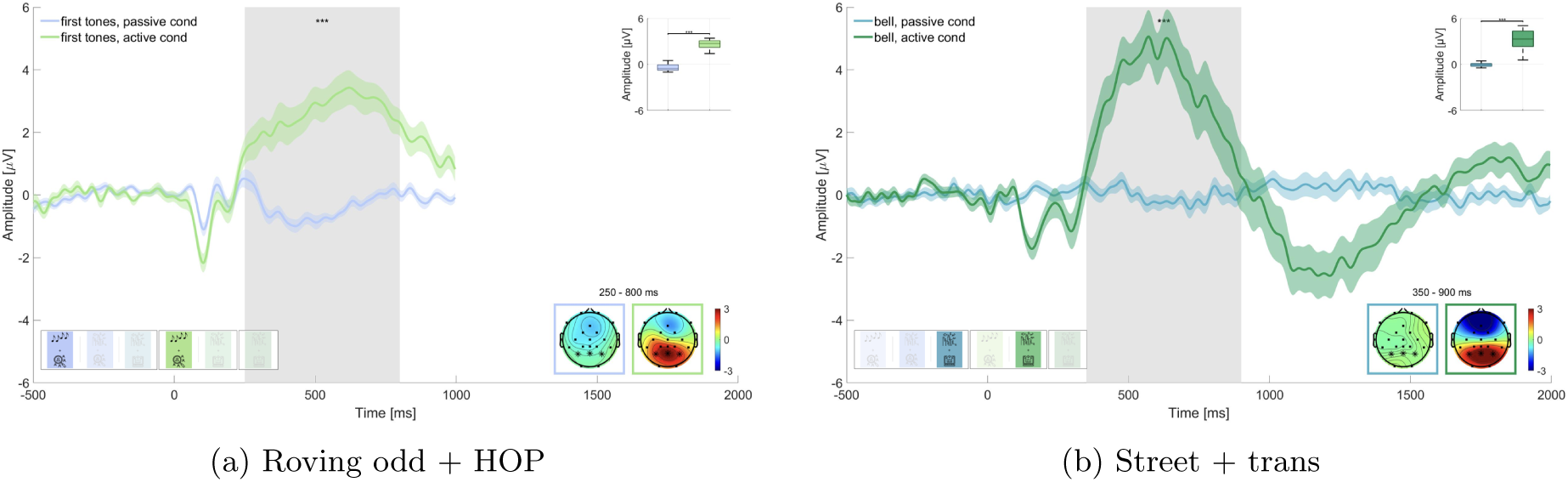
Comparison of the P300 amplitude for target sounds in passive vs active listening. Graphs show the average over the marked channels in the topographies. Asterisks indicate the level of significance (* p *≤* 0.05; ** p *≤* 0.01; *** p *≤* 0.001). **a** Roving oddball (roving odd) conditions. First tones in passive vs. active listening while performing the hidden object picture task (HOP). **b** Street conditions. Bell sounds in passive vs. active listening while performing the transcription task (trans).

For Hypothesis 2, we expected that irrelevant sounds in the same blocks as for hypothesis 1 (i.e., block 1 and 4: last tones and block 3 and 5: bird sounds) would elicit a larger N100 amplitude in the active listening than in the passive listening condition. Results are displayed in Figure 4. Wilcoxon signed-rank test revealed a significant difference for the roving oddball, where the amplitude of last tones was significantly larger in the active listening than in the passive listening condition (p *≤* 0.01, Z = 4.623, adjusted for 2 comparisons, Figure 4a). The mean amplitude in the time window of interest was m = -0.212 (SD = 1.100) in the passive listening condition and m = -0.830 (SD = 1.269) in the active listening condition. This was not found for the street soundscape, where bird sounds did not show a statistically significant difference between the two conditions (adj. p = 0.539, Z = 0.615, adjusted for 2 comparisons, Figure 4b). Here the amplitude in the time window of interest was m = 0.769 (SD = 0.920) in the passive listening condition and m = 0.647 (SD = 0.298) in the active listening condition. We found evidence in favor of hypothesis 2 concerning the effect in the roving oddball but not concerning the effect in the street soundscape.

**Figure 4:**
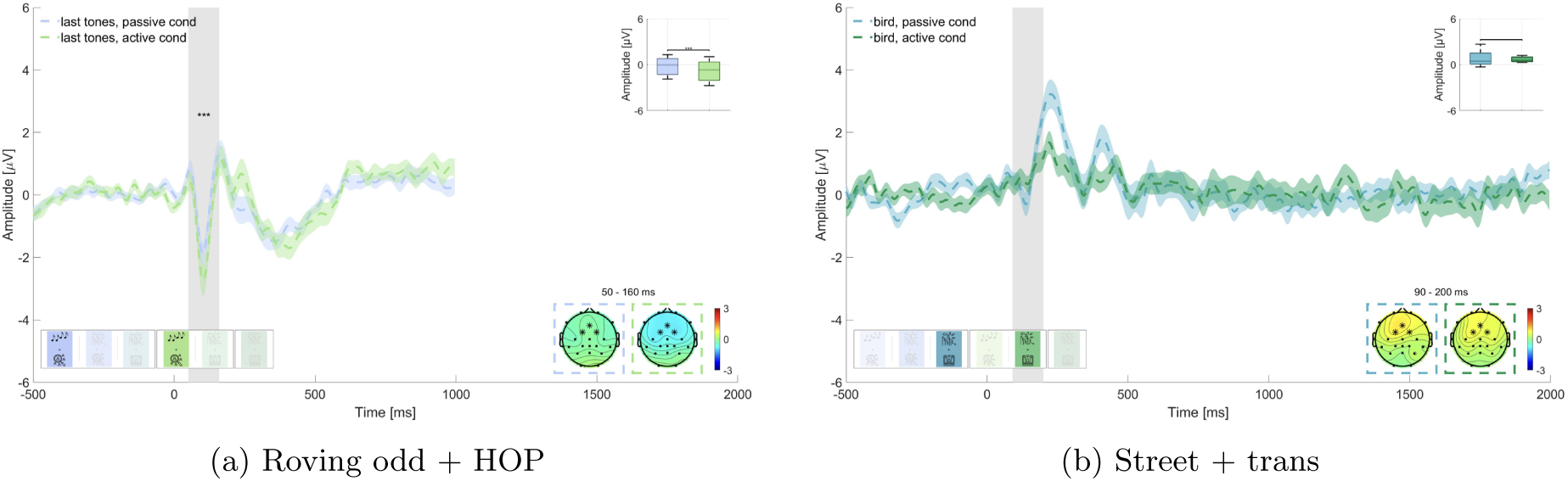
Comparison of the N100 amplitude for non-target sounds in passive vs active listening. Graphs show the average over the marked channels in the topographies. Asterisks indicate the level of significance (* p *≤* 0.05; ** p *≤* 0.01; *** p *≤* 0.001). **a** Roving oddball (roving odd) conditions. Last tones in passive vs. active listening while performing the hidden object picture task (HOP). **b** Street conditions. Bird sounds in passive vs. active listening while performing the transcription task (trans).

For Hypothesis 3 we were interested in the magnitude of the difference between P300 amplitudes in active and passive listening and expected a larger amplitude for the difference wave of the street soundscape conditions than for the roving oddball conditions. Therefore, we computed the difference waves for first tones in active listening (block 4) minus first tones in passive listening (block 1) and for bell sounds in active listening (block 5) minus bell sounds in passive listening (block 3). Wilcoxon signed-rank test revealed that the two waves were significantly different (adj. p *≤* 0.001, Z = -3.629, adjusted for 6 comparisons), but in the opposite direction than expected (see Figure 5). The difference wave for the street soundscape showed a larger amplitude (m = 3.329, SD = 1.341) than the difference wave for the roving oddball (m = 3.033, SD = 0.741). We therefore could not find evidence for hypothesis 3.

**Figure 5:**
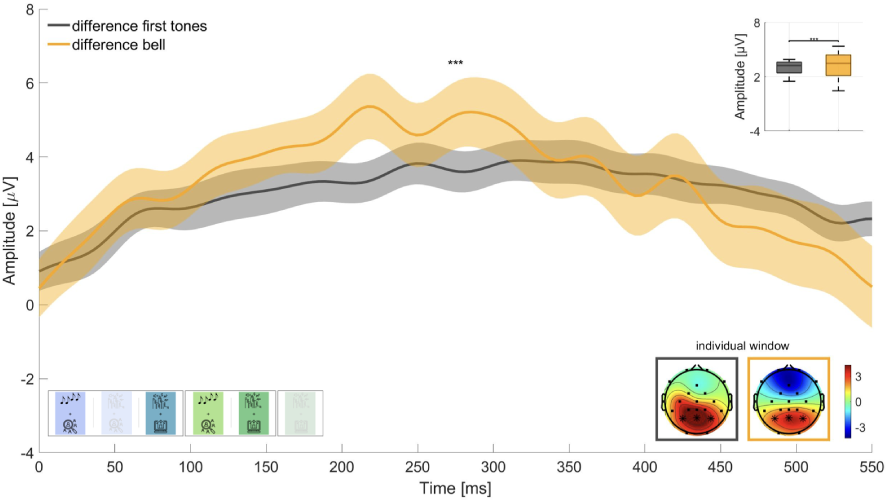
Difference waves for the P300 amplitude to target sounds in active minus passive listening. Graphs show the average over the marked channels in the topographies. Asterisks indicate the level of significance (* p *≤* 0.05; ** p *≤* 0.01; *** p *≤* 0.001).

In hypothesis 4 we investigated the effect of the additional task under passive listening, where we hypothesized to find smaller N100 amplitudes in the street soundscape condition while working on the transcription task (block 3), which is closer to everyday life, than while working on the more experimental hidden object picture task (block 2). We tested this for bell and bird sounds respectively (Figure 6). Wilcoxon signed-rank test revealed a statistically significant difference between the bird sounds while working on the hidden-object picture task and the transcription task (adj. p *≤* 0.001, Z = -4.031, adjusted for 6 comparisons, Figure 6a), where the amplitude on the N100 was significantly larger while working on the hidden-object picture task (m = 0.1852, SD = 0.760) than while working on the transcription task (m = 0.769, SD = 0.920). We found no statistically significant difference for the bell sounds (adj. p = 0.050, Z = -1.958, adjusted for 6 comparisons, Figure 6b) in the expected direction, where the bell sound while working on the hidden-object picture task had a larger N100 amplitude (m = -0.371, SD = 0.874) than while working on the transcription task (m = 0.029, SD = 0.410). We therefore found evidence in favor of Hypothesis 4 concerning the bird sounds but not the bell sounds.

**Figure 6:**
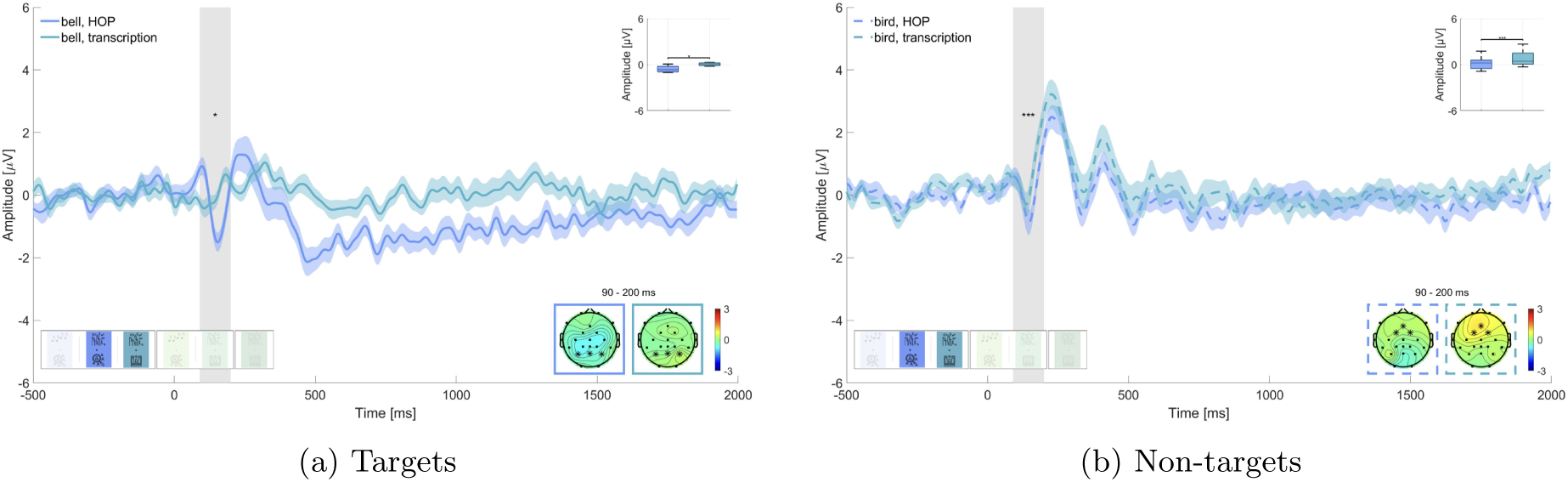
Comparison of the N100 amplitude in passive listening while either performing the hidden-object picture task or the transcription task. Graphs show the average over the marked channels in the topographies. Asterisks indicate the level of significance (* p *≤* 0.05; ** p *≤* 0.01; *** p *≤* 0.001). **a** Street conditions. Target sounds under the hidden-object picture task (HOP) and the transcription task. **b** Street conditions. Non-target sounds under the hidden-object picture task (HOP) and the transcription task.

In hypotheses 5 we tested whether we could effectively down-modulate the relevance of the bell in the wash-out block (block 6). We split the data into thirds and compared the P300 amplitude of the respective thirds of passive naive listening (block 3) and the wash-out block (block 6), as can be seen in Figure 7. We found that in the first third of the data the P300 amplitude of the wash-out block (m = 0.145, SD = 1.094) was significantly larger than in the passive naive listening block (m = -0.450, SD = 0.317; adj. p *≤* 0.01 Z = -4.851, adjusted for 6 comparisons, Figure 7a). In the second third the P300 amplitude of the wash-out block (m = 0.057, SD = 0.616) was not significantly different from the passive naive listening block (m = -0.108, SD = 0.310; adj. p = 0.022, Z = -2.288, adjusted for 6 comparisons, Figure 7b). For the final third, we found that, contrary to our expectation, the P300 amplitude in the passive naive listening condition was larger (m = 0.456, SD = 0.243) than in the wash-out condition (m = -0.631, SD = 0.284; adj. p *≤* 0.01, Z = 10.192, adjusted for 6 comparisons, Figure 7c).

**Figure 7:**
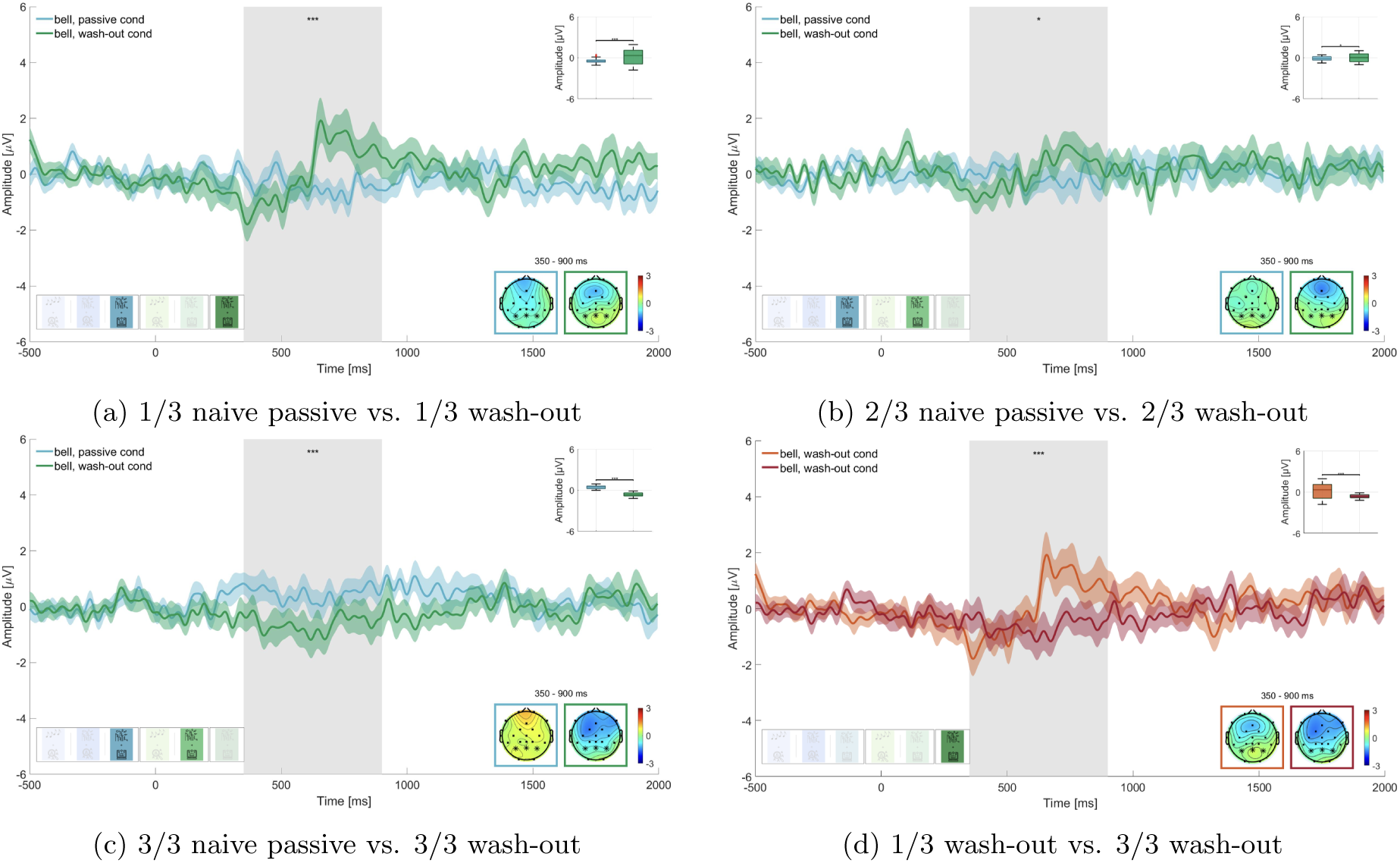
Comparison of the P300 amplitude in the passive naive listening condition vs wash-out condition. Graphs show the average over the marked channels in the topographies. Asterisks indicate the level of significance (* p *≤* 0.05; ** p *≤* 0.01; *** p *≤* 0.001). **a** Street conditions. First third of all bell sounds while working on the transcription task in naive passive listening vs. wash-out. **b** Street conditions. Second third of all bell sounds while working on the transcription task in naive passive listening vs. wash-out. **c** Street conditions. Third third of all bell sounds while working on the transcription task in naive passive listening vs. wash-out. **d** Street conditions. First third of all bell sounds while working on the transcription task in wash-out vs. third third of all bell sounds while working on the transcription task in wash-out.

To further investigate the wash-out effect, we also compared in the wash-out block the first third of trials to the last third of trials (Figure 7d) and found that the P300 amplitude was significantly larger in the first third of trials (m = 0.145, SD = 1.094) than in the final third of trials (m = -0.631, SD = 0.284; adj. p *≤* 0.01, Z = 6.672, adjusted for 4 comparisons).

## 4 Discussion

This study is part of a larger research project aimed at understanding the neural basis of sound perception in everyday life. Through a structured experimental design, we examined how to analyze EEG data in response to complex soundscapes within an office-like environment. By systematically increasing the complexity of the acoustic scene and the non-auditory task, our approach allows us to gain insights into how EEG data can be used to study (changes in) auditory processing in more ecologically valid conditions and over the course of more than 3 hours. This stepwise approach is essential for bridging the gap between the extensive body of lab-based research and the growing interest in understanding brain function in real-world scenarios.

### 4.1 Same Sound, Different Response - How Context Forms our Perception

Our study investigated the influence of contextual factors, particularly personal relevance, task complexity and stimuli properties, on sound perception. The results highlight how the properties of the non-auditory task significantly impact neural activity, specifically the N100 and P300 components. These findings emphasize that sound perception is shaped not only by the physical characteristics of stimuli but also by the context in which they are encountered, including the cognitive demands placed on the listener (Asutay & Västfjäll, 2012; Debnath & Wetzel, 2022; Rimskaya-Korsakova et al., 2022; Rosenkranz et al., 2023; Schlossmacher et al., 2021; Shinn-Cunningham & Best, 2008; Siegel & Stefanucci, 2011). This reinforces the idea that personal relevance and task complexity are critical factors in shaping sound perception, as introduced in our initial research questions.

#### 4.1.1 Stability of the P300 Component

Our results demonstrate that the P300 component is relatively stable across different auditory scenes (Hypothesis 1, Figure 3). The P300 exhibited a similar morphology in both the roving oddball and street soundscape conditions, suggesting that it is a robust measure of attention that is less influenced by the properties of the stimuli or the nature of the non-auditory task, which is in line with other research on the stability of the P300 component (Fallgatter et al., 2000). This finding reinforces the idea that the P300 can reliably indicate attentional processes in various contexts, making it a valuable tool in both laboratory and real-world settings (Polich, 2007). This stability across both isolated sounds and complex soundscapes suggests that the P300 as a marker of attention is a reliable measure across different auditory contexts, addressing our second research question.

However, contrary to our expectations, we observed a smaller amplitude difference in the P300 window between passive and active listening for the street soundscape compared to the HOP (Hypothesis 3, Figure 5). Although we hypothesized that the roving oddball would show a larger amplitude difference, the opposite was true. This unexpected finding could be attributed to the fact that the bell did not elicit a response within the designated time frame in the passive listening condition but a larger response in the active listening condition (cf. Figure 3b), leading to a larger difference in the N100 amplitude. With regard to the roving oddball, a slight potential was observed at the outset of the window of interest for the target sounds in the passive listening condition and a larger response in the active listening condition (cf. Figure 3a). This resulted in a smaller difference between passive and active listening, leading to the difference wave for the HOP being smaller than the difference wave for the street soundscape condition. As a limitation, the difference curve in the P300 window might not have aligned with our expectations due to the significant disparity in the number of trials analyzed. There were a maximum of 50 bell sounds in the street soundscape and up to 120 first tones in the HOP. The smaller number of trials in the street soundscape condition may have resulted in a larger neural response since the bell sound occurred only sporadically, as compared to the first tones in the HOP (Fitzgerald & Picton, 1984). Lastly, the low amount of trials in the street soundscape condition might have led to less reliable data, contributing to the unexpected outcomes in the difference wave analysis.

#### 4.1.2 Influence of Task Complexity on the N100 Component

In contrast to the P300, the N100 component appears to be more sensitive to the properties of the stimuli and the non-auditory task. While the N100 response was stable for the non-relevant last beep tones in the HOP condition, we did not observe a comparable N100 response to the non-relevant bird calls in the street soundscape (Hypothesis 2, Figure 4. Interestingly, there was an N100-like component in the passive listening condition, but this component was absent in the active listening condition. This discrepancy might indicate cognitive resource allocation, as proposed in the attentional resources hypothesis (Huang & Elhilali, 2020), where resources in the active condition were potentially used up by the transcription task and the detection of the behaviorally relevant bells. This is further supported by our finding that the non-auditory task had a strong effect on the N100 response to bell sounds (Hypothesis 4, Figure 6. The N100 was present only during the simpler hidden-object picture task and was completely absent during the more complex transcription task. This suggests that the more demanding task may have consumed the cognitive resources necessary for processing of the irrelevant bell sounds. The observed variability in the N100 response further indicates that complex soundscapes, especially when combined with more demanding tasks, lead to more context-dependent neural responses compared to isolated sounds.

We can rule out the possibility that the bell was simply not perceivable, as the same soundscape was played under both non-auditory tasks. Interestingly, the N100 response to the bird calls was observed under both tasks, although the amplitude was greater during the simpler task. This difference between the responses to the bell and bird calls might be due to the higher salience of the bird calls, which may have captured bottom-up attentional resources even when resources were otherwise allocated to the more complex task. An alternative explanation is that the hidden-object picture task always preceded the transcription task, which might have affected participants’ neural responsiveness, leading to a reduced N100 in the transcription task due to prior exposure or task-related fatigue. However, this explanation is not supported by the behavioral data of the non-auditory task, as there are no significant differences in performance that would indicate exhaustion of the participants. It is unlikely that the absence of an N100 component for the bell sounds while working on the transcription task is a mere order effect, since it is not seen for the bird sounds. As a limitation to our study design, we did not include a questionnaire on the awareness of the additional sounds in the street soundscape which in hindsight would have provided further validation of our findings through the use of an additional behavioral measure.

Overall, our findings demonstrate the complex interaction of task complexity and sound characteristics. A defining characteristic of natural sound is that it does not inherently possess a sharp sound onset. This is particularly crucial when examining stimuli that are not inherently salient and are embedded within the context of ambient sound. Ambient sound can result in the energetic masking of target stimuli, which in turn affects their perceptual onset (Oganian & Chang, 2019; Weise, Schröger, Fehér, Folyi, & Horváth, 2012). This may result in temporal smearing of the ERP, given that the perceptual onset is highly individual. Thus, when using more natural stimuli, researchers should be aware of the individual perceptual differences which may influence the effect. While more individualized analyses pipelines could be employed, e.g., by using individual time-windows to compute amplitude averages, our data showed a homogeneous perceptual onset. The high degree of coherence between participants is visible in the morphology of the ERP, where peaks are narrow.

#### 4.1.3 Learning and Unlearning Personal Relevance

Our study underlines, that relevance of sound can be learned and subsequently unlearned, independently of its physical properties. This is apparent from the gradual decrease in the P300 amplitude during the wash-out block (Hypothesis 5, Figure 7. Our findings with complex stimulus material in a simulated office environment are in line with previous studies (Polich, 2007) and add on the opposite effect, where relevant stimuli can be unlearned again. This suggests that relevance of sound can effectively be unlearned. Nevertheless, it remains unclear whether this is a consequence of the passage of time or an effect of the number of repetitions of the former target sounds. In order to investigate this point further, these two alternative explanations should be tested independently of each other in a future study.

Moreover, it would be intriguing to ascertain whether noise-sensitive individuals demonstrate a comparable pattern of unlearning relevance to non-noise-sensitive individuals in a comparative study. If noise-sensitive individuals exhibit greater difficulty in unlearning the relevance of a specific sound, this could serve as a potential avenue for developing interventions to address noise sensitivity and would extend the findings on a potentially impaired gating mechanism in noise-sensitive individuals (Shepherd, Hautus, Lee, & Mulgrew, 2016; Shepherd et al., 2019).

However, contrary to our expectations, the P300 amplitude in the final third of trials during the wash-out block was significantly lower than in the passive listening condition. While this could indicate an effect of intentional forgetting (Ten Oever, Sack, Oehrn, & Axmacher, 2021), it is more likely due to random data drift, given the small number of trials analyzed in the street soundscape condition after splitting into thirds. We had to balance two competing factors: maintaining a naturalistic soundscape and ensuring enough trials for analysis. The choice to include 50 target sounds over 45 minutes helped preserve the natural feel of working next to an open window in the city, but it may have compromised data quality, particularly for the split-into-thirds analysis.

### 4.2 Bridging Laboratory and Real-world Studies

Our study bridges the gap between controlled laboratory experiments and the unpredictable nature of real-world environments. By examining neural processing in a setting that mimics real-world conditions, we sought to understand how sound processing changes under different levels of experimental control. The results show that sound processing is influenced by multiple factors, including the complexity of the task, personal relevance, the properties of the sounds, and the broader environmental context. These findings suggest that certain neural markers, like the P300, generalize well from controlled lab environments to more complex, real-world-like scenarios. However, other components, such as the N100, exhibit greater sensitivity to task demands and may be less easily generalizable, thereby addressing our third research question.

We found that even small changes in the task setup, such as whether participants’ hands rested on the response key or had to move to it, led to significant differences in the recorded signal. We observed fewer artifacts in the hidden-object picture task, where there was a higher degree of experimental control, introduced by the fact that participants could keep their hand on the response key. In contrast, during the transcription task, which required more free movement of the hands and head (as participants looked down to locate keys), a substantial time-locked artifact emerged in the EEG data. These findings underscore how minor variations in task setup, particularly related to motor activity, can greatly impact data quality and should be carefully considered when interpreting results in less controlled environments (Gramann, 2024; Gramann et al., 2014; Jacobsen et al., 2021).

This highlights a general challenge when working with data of more naturalistic settings. While there are valid reasons to maintain consistent pre-processing across conditions, as we did following the protocol by Klug and Gramann (2021), this may not be ideal when artifact structures differ significantly between tasks. In our study, appending the data from all conditions before applying independent component analysis (ICA) ensured uniform pre-processing, allowing for cross-block comparisons. However, this approach may have resulted in the neglect of task-specific artifacts, which might have been more effectively reduced by a pre-processing method tailored to this specific task.

### 4.3 Implications for Future Research

Our findings demonstrate the feasibility of obtaining valuable insights from EEG data in realistic conditions, aligning with previous work in this area (Gramann et al., 2014; Jacobsen et al., 2021; Ladouce et al., 2019; Reiser et al., 2021; Rosenkranz, Haupt, Jaeger, Uslar, & Bleichner, 2024; Scanlon et al., 2022; Straetmans et al., 2021; Zink et al., 2016). Importantly, we showed that meaningful brain signals can be recorded over extended periods, even in the absence of specific instructions related to the sound stimuli. This opens new possibilities for studying sound perception in everyday life under natural conditions, without relying on momentary assessment. The ability to capture sound processing without active participant engagement is particularly promising for fields such as psychiatry, where it may help in studying impaired sensory gating in conditions like borderline personality disorder and schizophrenia, or in patients who may not comply with momentary assessment. The capacity to record EEG over extended periods in naturalistic settings, as demonstrated in our study, is pivotal for elucidating the neural mechanisms underlying the processing of complex auditory environments in everyday life. Importantly, our findings also underscore the necessity of preserving data integrity, as minor behavioral and environmental alterations can introduce unanticipated variability.

In light of these insights, future research should further explore sound perception in ecologically valid settings, while meticulously controlling for variables such as task complexity, environmental context, and personal relevance. Rather than a comprehensive transition to real-world experimentation, which introduces numerous uncontrolled variables, we propose a stepwise approach. This method permits researchers to incrementally introduce greater complexity while conducting validation checks between experimental blocks, thereby ensuring that observed neural changes are attributable to experimental manipulations rather than uncontrolled factors.

Specifically, future studies should examine how elements such as the acoustic properties of stimuli, task demands, and personal relevance shape neural processing. A more profound comprehension of these elements will empower researchers to devise experiments that capture real-world auditory complexity while maintaining scientific rigor.

The broader implications of our findings extend to other fields of beyond-the-lab research. As the number of studies investigating cognitive and neural processes in naturalistic environments increases, it is becoming increasingly evident that minor variations in task or environmental conditions can have a considerable impact on neural responses. This highlights the importance of meticulous experimental design and careful consideration of the context in which data is collected. By adopting a controlled, stepwise approach, researchers can systematically address the challenges of studying auditory perception in naturalistic environments, such as the variability introduced by task demands and environmental factors, which we highlighted in the introduction.

### 4.4 Conclusion: Insights on Sound Perception

In our study, we explored three key questions related to sound perception in naturalistic environments: (1) how personal relevance affects the perception of soundscapes over time, (2) how neural responses differ between isolated sounds and complex soundscapes, and (3) whether lab-based findings generalize to more naturalistic settings.

Our results suggest that sound perception is highly context-dependent, with task complexity and personal relevance playing significant roles in shaping neural responses, as mirrored by the N100 and P300 component. We demonstrated that the P300 is relatively stable across different auditory conditions, making it a reliable marker of attention even in dynamic environments. However, the N100 showed greater variability, suggesting that it is more sensitive to the complexity of the task and properties of the stimuli. Our findings further emphasize that personal relevance plays a critical role in determining how individuals process auditory stimuli in complex environments, further supporting the idea that sound perception is not only a function of the physical properties of sound but also its relevance to the listener.

We also found that results from controlled lab environments may not fully generalize to real-world settings, as task demands and environmental factors can significantly alter neural responses. However, by adopting a middle-ground approach, our study provides insights into how EEG can be used to study sound perception in more ecologically valid environments without sacrificing the precision of lab-based studies. Future research should continue to explore these questions, refining experimental designs to better capture the complexities of real-world auditory processing.

## 5 Data and Code Availability

The paradigm, analysis scripts and data to reproduce the findings of this paper can be found at: https://zenodo.org/records/14011457

## 6 Author Contributions

SK and MB conceptualized the experiment. SK performed the data acquisition, analyzed the data and wrote the manuscript to which MB, MJ and MR contributed with critical revisions. All authors approved the final version and agreed to be accountable for this work.

## 8 Acknowledgements

The authors would like to thank Negar Dadkhah and Amrah Gasimli for their valuable help with data collection and Johannes Freese who provided the bird sounds from his private data base. We would also like to thank the Friedrich Ebert Foundation, which always provides helpful support to SK.

## 10 Supplementary Material

### 10.1 Participant exclusion

### 10.2 Rationale for the multiple comparison correction

The following corrections were applied in each hypothesis:

- Hypothesis 1 involved two comparisons: First tones epochs block 1 vs. first tones epochs block 4 and bells epochs block 3 vs. bells epochs block 5. Since the bells epochs from block 3 were used in 6 comparisons across Hypotheses 1, 3, 4, and 5, we applied a Bonferroni correction for 6 comparisons to all tests within Hypothesis 1. This ensures that the repeated use of bells epochs from block 3 across multiple hypotheses is properly accounted for.
- Hypothesis 2 involved two comparisons: last tones epochs block 1 vs. last tones epochs block 4 and birds epochs block 3 vs. birds epochs block 5. Here, the birds epochs from block 3 were used in only 2 comparisons across Hypotheses 2 and 4. Therefore, a Bonferroni correction for 2 comparisons was applied to all tests within hypothesis 2.
- Hypothesis 3 involved one comparison: first tones epochs block 4 minus first tones epochs block 1 vs. bells epochs block 5 minus bells epochs block 3. Given that the bells epochs from block 3 were used in 6 comparisons across multiple hypotheses, the correction for this hypothesis was also based on 6 comparisons.
- Hypothesis 4 included two comparisons: bells epochs block 2 vs. bells epochs block 3 and birds epochs block 2 vs. birds epochs block 3. Since bells epochs from block 3 were used in 6 comparisons and birds epochs from block 3 in 2 comparisons across multiple hypotheses, we applied a Bonferroni correction for 6 comparisons, ensuring consistency within the hypothesis while accounting for the maximum number of relevant tests.
- Hypothesis 5 involved four comparisons between thirds of bell epochs from block 3 and Block 6. As bell epochs from block 3 were used in 6 comparisons across hypotheses, a Bonferroni correction for 6 comparisons was applied to all tests within Hypothesis 5.

## Footnotes

1 https://www.youtube.com/watch?v=Leg4s6KloU, (accessed 01.07.22)

2 https://www.youtube.com/watch?v=uya2CXwCY5w, (accessed 01.07.22)

3 for further information see: https://www.zooniverse.org/projects/courtaulddigital/world-architecture-unlocked/about/research

4 https://github.com/labstreaminglayer/liblsl-Matlab, v1.14.0.

5 https://github.com/labstreaminglayer/App-Input, v1.15.0.

6 https://github.com/labstreaminglayer/App-LabRecorder, v1.14.0.

## References

Asutay, E., & Västfjäll, D. (2012). Perception of Loudness Is Influenced by Emotion. PLoS ONE, 7 (6), e38660. doi: 10.1371/journal.pone.0038660

Basner, M., Babisch, W., Davis, A., Brink, M., Clark, C., Janssen, S., & Stansfeld, S. (2014). Auditory and non-auditory effects of noise on health. The Lancet, 383, 1325–1332. Retrieved from https://linkinghub.elsevier.com/retrieve/pii/S014067361361613X doi: 10.1016/S0140-6736(13)61613-X

Basner, M., Müller, U., & Elmenhorst, E.-M. (2011). Single and Combined Effects of Air, Road, and Rail Traffic Noise on Sleep and Recuperation. Sleep, 34 (1), 11–23. doi: 10.1093/sleep/34.1.11

Brainard, D. H. (1997). The Psychophysics Toolbox. Spatial Vision, 10 (4), 433–436. doi: 10.1163/156856897X00357

Brink, M., Omlin, S., Müller, C., Pieren, R., & Basner, M. (2011). An event-related analysis of awakening reactions due to nocturnal church bell noise. Science of The Total Environment, 409 (24), 5210–5220. doi: 10.1016/j.scitotenv.2011.09.020

Debnath, R., & Wetzel, N. (2022). Processing of task-irrelevant sounds during typical everyday activities in children. Developmental Psychobiology, 64 (7), e22331. doi: 10.1002/dev.22331

Dehais, F., Roy, R., & Scannella, S. (2019). Inattentional deafness to auditory alarms: Inter-individual differences, electrophysiological signature and single trial classification. Behavioural Brain Research, 360, 51–59. doi: 10.1016/j.bbr.2018.11.045

Delorme, A., & Makeig, S. (2004). EEGLAB: An open source toolbox for analysis of single-trial EEG dynamics including independent component analysis. Journal of Neuroscience Methods, 134 (1), 9–21. doi: 10.1016/j.jneumeth.2003.10.009

Deshpande, P., Brandt, C., Debener, S., & Neher, T. (2022). Comparing Clinically Applicable Behavioral and Electrophysiological Measures of Speech Detection, Discrimination, and Comprehension. Trends in Hearing, 26 , 233121652211397. doi: 10.1177/23312165221139733

Ding, N., & Simon, J. Z. (2012). Emergence of neural encoding of auditory objects while listening to competing speakers. Proceedings of the National Academy of Sciences, 109 (29), 11854–11859. doi: 10.1073/pnas.1205381109

Fallgatter, A. J., Eisenack, S. S., Neuhauser, B., Aranda, D., Scheuerpflug, P., & Herrmann, M. J. (2000). Stability of Late Event-Related Potentials: Topographical Descriptors of Motor Control Compared with the P300 Amplitude. Brain Topography, 12 (4), 255–261. doi: 10.1023/A:1023403420864

Fitzgerald, P. G., & Picton, T. W. (1984). The Effects of Probability and Discriminability on the Evoked Potentials to Unpredictable Stimuli. Annals of the New York Academy of Sciences, 425 (1), 199–203. doi: 10.1111/j.1749-6632.1984.tb23533.x

Fredrickson, B. L., & Kahneman, D. (1993). Duration neglect in retrospective evaluations of affective episodes. Journal of Personality and Social Psychology, 65 (1), 45–55. doi: 10.1037/0022-3514.65.1.45

Fuglsang, S. A., Dau, T., & Hjortkjær, J. (2017). Noise-robust cortical tracking of attended speech in real-world acoustic scenes. NeuroImage, 156, 435–444. doi: 10.1016/j.neuroimage.2017.04.026

Garrido, M. I., Friston, K. J., Kiebel, S. J., Stephan, K. E., Baldeweg, T., & Kilner, J. M. (2008). The functional anatomy of the MMN: A DCM study of the roving paradigm. NeuroImage, 42 (2), 936–944. doi: 10.1016/j.neuroimage.2008.05.018

Gorgolewski, K. J., Auer, T., Calhoun, V. D., Craddock, R. C., Das, S., Duff, E. P., … Poldrack, R. A. (2016). The brain imaging data structure, a format for organizing and describing outputs of neuroimaging experiments. Scientific Data, 3 (1), 160044. doi: 10.1038/sdata.2016.44

Gramann, K. (2024). Mobile EEG for Neurourbanism Research - What Could Possibly Go Wrong? A Critical Review with Guidelines. doi: 10.1101/2024.03.22.586309

Gramann, K., Ferris, D. P., Gwin, J., & Makeig, S. (2014). Imaging natural cognition in action. International Journal of Psychophysiology, 91 (1), 22–29. doi: 10.1016/j.ijpsycho.2013.09.003

Hasson, U., & Honey, C. J. (2012). Future trends in Neuroimaging: Neural processes as expressed within real-life contexts. NeuroImage, 62 (2), 1272–1278. doi: 10.1016/j.neuroimage.2012.02.004

Heinonen-Guzejev, M. (2008). Noise sensitivity: Medical, psychological and genetic aspects. Helsinki: University of Helsinki.

Hjortskov, M. (2017). Priming and context effects in citizen satisfaction surveys. Public Administration, 95 (4), 912–926. doi: 10.1111/padm.12346

Hölle, D., & Bleichner, M. G. (2023). Smartphone-based ear-electroencephalography to study sound processing in everyday life. European Journal of Neuroscience, 58 (7), 3671–3685. doi: 10.1111/ejn.16124

Hölle, D., Meekes, J., & Bleichner, M. G. (2021). Mobile ear-EEG to study auditory attention in everyday life: Auditory attention in everyday life. Behavior Research Methods, 53 (5), 2025– 2036. doi: 10.3758/s13428-021-01538-0

Holtze, B., Jaeger, M., Debener, S., Adiloğlu, K., & Mirkovic, B. (2021). Are They Calling My Name? Attention Capture Is Reflected in the Neural Tracking of Attended and Ignored Speech. Frontiers in Neuroscience, 15, 643705. doi: 10.3389/fnins.2021.643705

Hong, J. Y., & Jeon, J. Y. (2015). Influence of urban contexts on soundscape perceptions: A structural equation modeling approach. Landscape and Urban Planning, 141, 78–87. doi: 10.1016/j.landurbplan.2015.05.004

Horton, C., Srinivasan, R., & D’Zmura, M. (2014). Envelope responses in single-trial EEG indicate attended speaker in a ‘cocktail party’. Journal of Neural Engineering, 11 (4), 046015. doi: 10.1088/1741-2560/11/4/046015

Huang, N., & Elhilali, M. (2020). Push-pull competition between bottom-up and top-down auditory attention to natural soundscapes. eLife, 9. doi: 10.7554/eLife.52984

Hygge, S., Evans, G. W., & Bullinger, M. (2002). A Prospective Study of Some Effects of Aircraft Noise on Cognitive Performance in Schoolchildren. Psychological Science, 13 (5), 469–474. doi: 10.1111/1467-9280.00483

Jacobsen, N. S. J., Blum, S., Witt, K., & Debener, S. (2021). A walk in the park? Characterizing gait-related artifacts in mobile EEG recordings. European Journal of Neuroscience, 54 (12), 8421–8440. doi: 10.1111/ejn.14965

Jaeger, M., Mirkovic, B., Bleichner, M. G., & Debener, S. (2020). Decoding the Attended Speaker From EEG Using Adaptive Evaluation Intervals Captures Fluctuations in Attentional Listening. Frontiers in Neuroscience, 14, 603. doi: 10.3389/fnins.2020.00603

Kjellberg, A., Landström, U., Tesarz, M., Söderberg, L., & Akerlund, E. (1996). The Effects of nonphysical noise characteristics, ongoing task and noise sensitivity on annoyance and distraction due to noise at work. Journal of Environmental Psychology, 16 (2), 123–136. doi: 10.1006/jevp.1996.0010

Kleiner, M., Brainard, D. H., & Pelli, D. G. (2007). “what’s new in psychtoolbox-3?” perception 36 ecvp abstract supplement.

Klug, M., & Gramann, K. (2021). Identifying key factors for improving ICA-based decomposition of EEG data in mobile and stationary experiments. European Journal of Neuroscience, 54 (12), 8406–8420. doi: 10.1111/ejn.14992

Krugliak, A., & Clarke, A. (2022, February). Towards real-world neuroscience using mobile EEG and augmented reality. Scientific Reports, 12 (1), 2291. doi: 10.1038/s41598-022-06296-3

Ladouce, S., Donaldson, D. I., Dudchenko, P. A., & Ietswaart, M. (2019). Mobile eeg identifies the re-allocation of attention during real-world activity. Scientific Reports, 9, 15851. doi: 10.1038/s41598-019-51996-y

Lorenzi, C., Apoux, F., Grinfeder, E., Krause, B., Miller-Viacava, N., & Sueur, J. (2023). Human Auditory Ecology: Extending Hearing Research to the Perception of Natural Soundscapes by Humans in Rapidly Changing Environments. Trends in Hearing, 27, 23312165231212032. doi: 10.1177/23312165231212032

Maselli, A., Gordon, J., Eluchans, M., Lancia, G. L., Thiery, T., Moretti, R., … Pezzulo, G. (2023). Beyond simple laboratory studies: Developing sophisticated models to study rich behavior. Physics of Life Reviews, 46, 220–244. doi: 10.1016/j.plrev.2023.07.006

Mathias, S. V., & Bensalem-Owen, M. (2019). Artifacts That Can Be Misinterpreted as Interictal Discharges. Journal of Clinical Neurophysiology, 36 (4), 264. doi: 10.1097/WNP.0000000000000605

Mehrotra, A., Shukla, S. P., Shukla, A., Manar, M. K., Singh, S., & Mehrotra, M. (2024). A Comprehensive Review of Auditory and Non-Auditory Effects of Noise on Human Health. Noise and Health, 26 (121), 59–69. doi: 10.4103/nah.nah_124_23

Mirkovic, B., Debener, S., Jaeger, M., & De Vos, M. (2015). Decoding the attended speech stream with multi-channel EEG: Implications for online, daily-life applications. Journal of Neural Engineering, 12 (4), 046007. doi: 10.1088/1741-2560/12/4/046007

Näätänen, R., Kujala, T., & Winkler, I. (2011). Auditory processing that leads to conscious perception: A unique window to central auditory processing opened by the mismatch negativity and related responses. Psychophysiology, 48 (1), 4–22. doi: 10.1111/j.1469-8986.2010.01114.x

Näätänen, R., & Picton, T. (1987). The N1 Wave of the Human Electric and Magnetic Response to Sound: A Review and an Analysis of the Component Structure. Psychophysiology, 24 (4), 375–425. doi: 10.1111/j.1469-8986.1987.tb00311.x

Nastase, S. A., Goldstein, A., & Hasson, U. (2020). Keep it real: Rethinking the primacy of experimental control in cognitive neuroscience. NeuroImage, 222, 117254. doi: 10.1016/j.neuroimage.2020.117254

O’Connell, A., & Greene, C. M. (2017). Not strange but not true: Self-reported interest in a topic increases false memory. Memory, 25 (8), 969–977. doi: 10.1080/09658211.2016.1237655

Oganian, Y., & Chang, E. F. (2019). A speech envelope landmark for syllable encoding in human superior temporal gyrus. Science Advances, 5 (11), eaay6279. doi: 10.1126/sciadv.aay6279

O’Sullivan, J. A., Power, A. J., Mesgarani, N., Rajaram, S., Foxe, J. J., Shinn-Cunningham, B. G., … Lalor, E. C. (2015). Attentional Selection in a Cocktail Party Environment Can Be Decoded from Single-Trial EEG. Cerebral Cortex, 25 (7), 1697–1706. doi: 10.1093/cercor/bht355

Paunović, K., Jakovljević, B., & Belojević, G. (2009). Predictors of noise annoyance in noisy and quiet urban streets. Science of The Total Environment, 407, 3707–3711. doi: 10.1016/j.scitotenv.2009.02.033

Pelli, D. G. (1997). The VideoToolbox software for visual psychophysics: Transforming numbers into movies. Spatial Vision, 10 (4), 437–442. doi: 10.1163/156856897X00366

Peris, E., Blane, N., Fons, J., Sainz de la Maza, M., José Ramos, M., Domingues, F., … Adams, M. (2020). Environmental noise in Europe, 2020. Luxembourg: Publications Office of the European Union.

Pernet, C. R., Appelhoff, S., Gorgolewski, K. J., Flandin, G., Phillips, C., Delorme, A., & Oostenveld, R. (2019). EEG-BIDS, an extension to the brain imaging data structure for electroencephalography. Scientific Data, 6 (1), 103. doi: 10.1038/s41597-019-0104-8

Pierrette, M., Marquis-Favre, C., Morel, J., Rioux, L., Vallet, M., Viollon, S., & Moch, A. (2012). Noise annoyance from industrial and road traffic combined noises: A survey and a total annoyance model comparison. Journal of Environmental Psychology, 32 (2), 178–186. doi: 10.1016/j.jenvp.2012.01.006

Pion-Tonachini, L., Kreutz-Delgado, K., & Makeig, S. (2019). ICLabel: An automated electroencephalographic independent component classifier, dataset, and website. NeuroImage, 198, 181–197. doi: 10.1016/j.neuroimage.2019.05.026

Polich, J. (2007). Updating P300: An integrative theory of P3a and P3b. Clinical Neurophysiology, 118 (10), 2128–2148. Retrieved from https://linkinghub.elsevier.com/retrieve/pii/S1388245707001897 doi: 10.1016/j.clinph.2007.04.019

Reiser, J. E., Wascher, E., Rinkenauer, G., & Arnau, S. (2021). Cognitive-motor interference in the wild: Assessing the effects of movement complexity on task switching using mobile EEG. European Journal of Neuroscience, 54 (12), 8175–8195. doi: 10.1111/ejn.14959

Rimskaya-Korsakova, L. K., Pyatakov, P. A., & Shulyapov, S. A. (2022). Evaluations of the Annoyance Effects of Noise. Acoustical Physics, 68 (5), 502–512. doi: 10.1134/S1063771022050098

Rosenkranz, M., Cetin, T., Uslar, V. N., & Bleichner, M. G. (2023). Investigating the attentional focus to workplace-related soundscapes in a complex audio-visual-motor task using EEG. Frontiers in Neuroergonomics, 3, 1062227. doi: 10.3389/fnrgo.2022.1062227

Rosenkranz, M., Haupt, T., Jaeger, M., Uslar, V. N., & Bleichner, M. G. (2024). Sound perception in realistic surgery scenarios: Towards EEG-based auditory work strain measures for medical personnel. doi: 10.1101/2024.05.07.592873

Roye, A., Jacobsen, T., & Schröger, E. (2013). Discrimination of personally significant from nonsignificant sounds: A training study. Cognitive, Affective, & Behavioral Neuroscience, 13 (4), 930–943. doi: 10.3758/s13415-013-0173-7

Scanlon, J., Jacobsen, N., Maack, M., & Debener, S. (2022). Stepping in time: Alpha-mu and beta oscillations during a walking synchronization task. NeuroImage, 253, 119099. doi: 10.1016/j.neuroimage.2022.119099

Schinkel-Bielefeld, N., Burke, L., Holube, I., Iankilevitch, M., Jenstad, L. M., Lelic, D., … Wu, Y.-H. (2024). Implementing Ecological Momentary Assessment in Audiological Research: Opportunities and Challenges. American Journal of Audiology, 33 (3), 648–673. doi: 10.1044/2024_AJA-23-00249

Schlossmacher, I., Dellert, T., Bruchmann, M., & Straube, T. (2021). Dissociating neural correlates of consciousness and task relevance during auditory processing. NeuroImage, 228, 117712. doi: 10.1016/j.neuroimage.2020.117712

Shepherd, D., Hautus, M. J., Lee, S. Y., & Mulgrew, J. (2016). Electrophysiological approaches to noise sensitivity. Journal of Clinical and Experimental Neuropsychology, 38 (8), 900–912. doi: 10.1080/13803395.2016.1176995

Shepherd, D., Lodhia, V., & Hautus, M. J. (2019). Electrophysiological indices of amplitude modulated sounds and sensitivity to noise. International Journal of Psychophysiology, 139, 59–67. Retrieved from https://linkinghub.elsevier.com/retrieve/pii/S0167876018309905 doi: 10.1016/j.ijpsycho.2019.03.005

Shinn-Cunningham, B. G., & Best, V. (2008). Selective Attention in Normal and Impaired Hearing. Trends in Amplification, 12 (4), 283–299. doi: 10.1177/1084713808325306

Siegel, E. H., & Stefanucci, J. K. (2011). A little bit louder now: Negative affect increases perceived loudness. Emotion, 11 (4), 1006–1011. doi: 10.1037/a0024590

Sonnleitner, A., Treder, M. S., Simon, M., Willmann, S., Ewald, A., Buchner, A., & Schrauf, M. (2014). EEG alpha spindles and prolonged brake reaction times during auditory distraction in an on-road driving study. Accident Analysis & Prevention, 62, 110–118. doi: 10.1016/j.aap.2013.08.026

Straetmans, L., Holtze, B., Debener, S., Jaeger, M., & Mirkovic, B. (2021). Neural tracking to go: Auditory attention decoding and saliency detection with mobile EEG. Journal of Neural Engineering, 18 (6), 066054. doi: 10.1088/1741-2552/ac42b5

Sumner, R. (2022). RovingMMN. Retrieved from https://github.com/RachaelSumner/RovingMMN

Ten Oever, S., Sack, A. T., Oehrn, C. R., & Axmacher, N. (2021). An engram of intentionally forgotten information. Nature Communications, 12 (1), 6443. doi: 10.1038/s41467-021-26713-x

Weinstein, N. D. (1978). Individual differences in reactions to noise: A longitudinal study in a college dormitory. Journal of Applied Psychology, 63 (4), 458–466. doi: 10.1037/0021-9010.63.4.458

Weise, A., Schröger, E., Fehér, B., Folyi, T., & Horváth, J. (2012). Auditory event-related potentials reflect dedicated change detection activity for higher-order acoustic transitions. Biological Psychology, 91 (1), 142–149. doi: 10.1016/j.biopsycho.2012.06.001

World Health Organization (Ed.). (2011). Burden of disease from environmental noise: Quantification of healthy life years lost in Europe. Copenhagen: World Health Organization, Regional Office for Europe.

Yong Jeon, J., Jik Lee, P., Young Hong, J., & Cabrera, D. (2011). Non-auditory factors affecting urban soundscape evaluation. The Journal of the Acoustical Society of America, 130 (6), 3761–3770. doi: 10.1121/1.3652902

Yoon, J.-H., Won, J.-U., Lee, W., Jung, P. K., & Roh, J. (2014). Occupational Noise Annoyance Linked to Depressive Symptoms and Suicidal Ideation: A Result from Nationwide Survey of Korea. PLoS ONE, 9 (8), e105321. doi: 10.1371/journal.pone.0105321

Zink, R., Hunyadi, B., Huffel, S. V., & Vos, M. D. (2016). Mobile EEG on the bike: Disentangling attentional and physical contributions to auditory attention tasks. Journal of Neural Engineering, 13 (4), 046017. doi: 10.1088/1741-2560/13/4/046017

